# 2-Methoxyestradiol Treatment Prevents Graft-versus-Host Disease While Preserving Graft-versus-Leukemia Effect in Mice

**DOI:** 10.1101/2025.08.11.667504

**Authors:** Dan Li, Roland Bassett, Richard E. Champlin, Jeffrey J. Molldrem, Tak W. Mak, Qing Ma

## Abstract

2-Methoxyestradiol (2ME2, Panzem) is an endogenous metabolite that is well-tolerated in phase I/II clinical trials for variety of tumors. The plasma levels of 2ME2 may increase up to 1,000-fold during pregnancy and correlate temporally with the remission of rheumatoid arthritis (RA) and multiple sclerosis (MS) symptoms. The anti-inflammatory properties of 2ME2 were recently established in the mouse model of MS, and the mechanism of action is the ability of 2ME2 to inhibit lymphocyte proliferation, cytokine production and T cell polarization. Herein, we have demonstrated that 2ME2 treatment can significantly reduce the mortality and morbidity associated with graft-versus-host disease (GVHD). There is a lower number of donor-derived CD4^+^ and CD8^+^ T cells in the peripheral lymph node and Peyer’s patches of 2ME2-treated mice compared to control recipients. Moreover, 2ME2 exposure can significantly decrease the production of IFN-γ and IL-2 in donor-derived CD4^+^ T cells and serum in GVHD mice. However, 2ME2 treatment has no effect on the differentiation of CD8^+^ effector T cells in vivo and their cytolytic activity remains intact. Furthermore, 2ME2 therapy is effective in preventing GVHD while preserving graft-versus leukemia (GVL) activity in mice. Our findings indicate that 2ME2 could be a novel and effective treatment for GVHD patients.

## INTRODUCTION

Allogeneic bone marrow transplantation (allo-HSCT) is an effective therapy for hematological malignancies^1-2^. But the limiting factor is GVHD, a result of alloimmune responses mediated by donor T lymphocytes and recipient antigen presenting cells (APCs)^2-4^. Pharmacologic prophylaxis has lowered the risk for acute GVHD, but still 35-50% of patients develop Grade II-IV acute GVHD^4-5^. For the past three decades steroids have been the standard initial therapy for grades II-IV acute GVHD. Various agents have been studied for the treatment of steroid-refractory acute GVHD with published response rates approximating 40%. Yet, even when patients responded, the outcome for patients with steroid-refractory acute GVHD is poor with a mortality rate of approximately 70%^4-5^. GVHD is caused by alloreactive donor T lymphocytes, which results in multi-organ dysfunction and destruction^1-3^. The main effector in GVHD are donor T cell activation, followed by proliferation and differentiation into activated effector cells, and finally specific tissue damage^3, 6-10^.

2-Methoxyestradiol (2ME2, Panzem) is an endogenous metabolite that at pharmacologic doses exerts antimitotic and anti-angiogenic properties^11-12^. 2ME2 can bind to tubulin and disrupt the microtubule dynamics, leading to the depolymerization of tubulin. As a result, rapidly dividing cells are undergoing G2/M arrest and apoptosis in the presence of 2ME2. 2ME2 has been well-tolerated and safe in phase I and II clinical trials targeting various tumor types^13^. The sharp increase of 2ME2 level in plasma during pregnancy has been shown to correlate with the temporary remission of multiple sclerosis (MS) and rheumatoid arthritis (RA) symptoms in pregnant women^4-17^. 2ME2 has also been shown to exhibit disease-modifying activity in autoimmune models of RA in rodents^18-20^. A deeper understanding of the anti-inflammatory properties of 2ME2 was recently established in mouse experimental autoimmune encephalitis (EAE), a rodent model of MS^21^. In addition to regulating lymphocyte activation and cytokine production as the mechanism of action, 2ME2 also inhibits IL-17 production in Th17-polarized cells in the pathogenesis of EAE^21-22^.

These above cited mouse autoimmune disease models serve as important surrogate models for GVHD, which was our impetus for further study of this compound. We demonstrated that 2ME2 treatment can reduce the mortality and morbidity associated with mouse GVHD. There was significantly reduced number of donor-derived CD4^+^ and CD8^+^ T cells in the secondary lymphoid organs of 2ME2-treated recipient mice. In additional studies, 2ME2 exposure was found to attenuate cytokine production, resulting in the reduced production of IFN-γ (Th1) and IL-2 in donor-derived CD4^+^ T cells and serum. Furthermore, 2ME2 treatment has no effect on the differentiation of CD8^+^ effector T cells and their cytolytic activity in vitro, and GVL effect remains intact in mice.

## MATERIALS AND METHODS

### Animals and Reagents

C57BL/6 (B6; H-2^b^) and BALB/c (H-2^d^) mice were purchased from the Jackson Laboratory. 2ME2 was provided by EntreMed, and stored and used as recommended according to the instruction from the manufacturer. The animal experiments were approved by the Institutional Animal Care and Use Committee at University of Texas M.D. Anderson Cancer Center.

### GVHD Induction and Pathology

The MHC class I and II disparate model, BALB/C (H-2^d^) to C57BL/6 (H-2^b^), was used to establish GVHD^23^. All recipients were age-matched females and 2-6 months of age at the time of BMT. To generate BMT chimeras, recipient Balb/C mice were irradiated with 8.5 Gy TBI (^137^Cs source) split into 2 doses at day -1, and received 5×10^6^ bone marrow (BM) cells plus 10×10^6^ splenocytes from donor C57BL/6 mice on day 0. Mice were weighed twice a week while survival and clinical signs of GVHD (hair loss, hunched back and diarrhea) were monitored daily. On day 7 post-transplant, GVHD target tissues including skin, liver, intestine, and lung were collected and fixed in 10% formalin for histopathological analysis.

### Analysis of Donor-derived T Cells

Recipient mice were sacrificed 7 days post-transplant. Spleen, lymph nodes and Peyer’s patches were harvested, and single cell suspension were stained with H-2D^b^ antibody to identify donor-derived cells in secondary lymph organs. Donor-derived T cells (H-2D^b+^), CD4^+^H-2D^b+^ and CD8^+^H-2D^b+^ subsets were collected and analyzed by FACS. The total number of each subset recovered from tissues was determined by Flowjo software.

### Mixed Lymphocyte Culture

The experiment was performed in 96-well microtiter plates. Responder T cells from C57BL/6 mice were plated at 1×10^6^ cells/ml in a volume of 200 µl/well and co-cultured at a ratio of 2:1 with 3400 cGy irradiated stimulator cells from Balb/C mice. For proliferation assay, responder cells were labeled with CFSE, and cell division was monitored by using the FITC channel in a flow cytometer as previously described^23^. For cell cycle analysis, responder cells were stained with Ki-67 antibody and PI. FACS was used to analyze and determine the G0, G1 and G2/M phases of the cell cycle as previously described^24^.

### Cytokine Flow Cytometry Assay

Intracellular cytokine production was measured using flow cytometry as previously described^25^. Briefly, one hour after simulation of T cells harvest from MLR or recipient mice, Brefeldin A was added to enable accumulation of intracellular cytokines. Following an additional 5 hours of incubation, cells were fixed and permeabilized for assessing the simultaneous expression of surface markers and intracellular cytokines. IL-2-, IL-17- and IFN-γ-producing cells were identified using intracellular staining, and donor-derived T cells were determined using CD4 and H-2D^b^ antibodies by FACS. The percentage of IL-2^+^, IL-17^+^ and IFNγ^+^ cells was analyzed on a LSR-II cytometer and determined by Flowjo software. Gates defining cytokine-positive populations were defined based on the upper limits of fluorescence of unstimulated cells stained with the same antibodies.

### Cytotoxicity Assay

The effector cells were activated C57BL/6 T cells generated from MLR with allo-stimulation for 5 days in vitro. The cytotoxic specificity of mouse cytolytic T lymphocytes (CTLs) was determined using the flow cytometry-based CTL assay measuring the cleavage of caspase-3 in targets cells as previously described^26^. The target cells were P815 labeled with a dye, and then incubated with CTLs at different effector:target (E:T) ratios for 3 to 4 hours and fixed/permeabilized, and stained with a anti-cleaved caspase-3 rabbit mAb. The stained cells were analyzed on FACS within 24 hrs.

### GVL Mouse Model and Imaging Analysis

P815 cells with a lentivirus vector coding for firefly luciferase (FLuc) for bioluminescence imaging were used for GVL model. Recipient BALB/c mice received P815-Fluc mastocytoma cells (3000) on the day of transplantation in addition to donor bone marrow with or without T cells. P815 growth and GVL effect were measured by bioluminescence imaging and emitted photons over time as described^27^. Briefly, mice were injected intraperitoneally with luciferin (10 μg/g body weight). After 10 minutes, mice were imaged using an IVIS200 charge-coupled device (CCD) imaging system (Xenogen, Alameda, CA) for 30 seconds to 5 minutes. The IVIS200 CCD imaging system was used to perform serial imaging studies twice a week post-transplant. Leukemia expansion was quantified in photons/second/cm^2^. Imaging data were analyzed and quantified with Living Image Software (Xenogen) and IgorProCarbon (WaveMetrics, Lake Oswego, OR). The photon flux (photons/second/cm^2^, photons over total body area) measurements were used to estimate the engraftment of P815-FLuc cells.

### Statistical Analysis

Survival data were plotted using the Kaplan-Meier method and analyzed by the log-rank test. A *p* value of 0.05 or less was considered statistically significant.

## RESULTS

### Reduced GVHD mortality and morbidity in 2ME2-treated recipient mice

To determine whether 2ME2 affects acute GVHD in mice, we used the well-established C57BL/6 (H-2^b^) into BALB/c (H-2^d^) MHC class I and II disparate model^23^. In the pilot dose finding study, mice were treated 24 hrs post-transplant with 2ME2 daily at 25, 50, 100, 150 and 300 mg/kg (i.p.). We found that 2ME2 treatment at 25 mg/kg dose can protect mice from GVHD related mortality (data not shown), and confirmed the previous data from the manufacture (EntreMed) that the tolerability to 2ME2 was much reduced in irradiated animals, and 150mg/kg was not tolerable (data not shown). In comparison to PBS control, 2ME2-treated mice showed a significant lower mortality rate (Figure 1A). Within 4 weeks post-transplantation, about 80% of 2ME2-treated recipients survived, compared with only about 20% of PBS control mice (P=0.004; n=15 in each group). Mice received BM only did not show any signs of GVHD (data not shown). On day 7 post-transplant, mice were sacrificed; their skin, small intestine, liver and lung were collected for histopathological analysis. As shown in Figure 1B, significantly increased inflammatory cells infiltration and apoptotic cells were presented in hair follicles of the skin. There were necrosis and apoptosis in the crypts of the small intestine; lymphoplasmacytic infiltration in portal tracts and necrosis of hepatocytes in the liver. Inflammatory infiltration were seen in thickened arterial and alveolar wall in the lung of PBS control mice. Whereas control recipients had severe GVHD in the skin, intestine, liver, and lung, 2ME2-treated mice exhibited only mild changes in these organs, reflected in their significantly lower GVHD scores (Figure 1C). Thus, 2ME2 treatment can reduce GVHD mortality and morbidity in mice.

**Figure 1.**
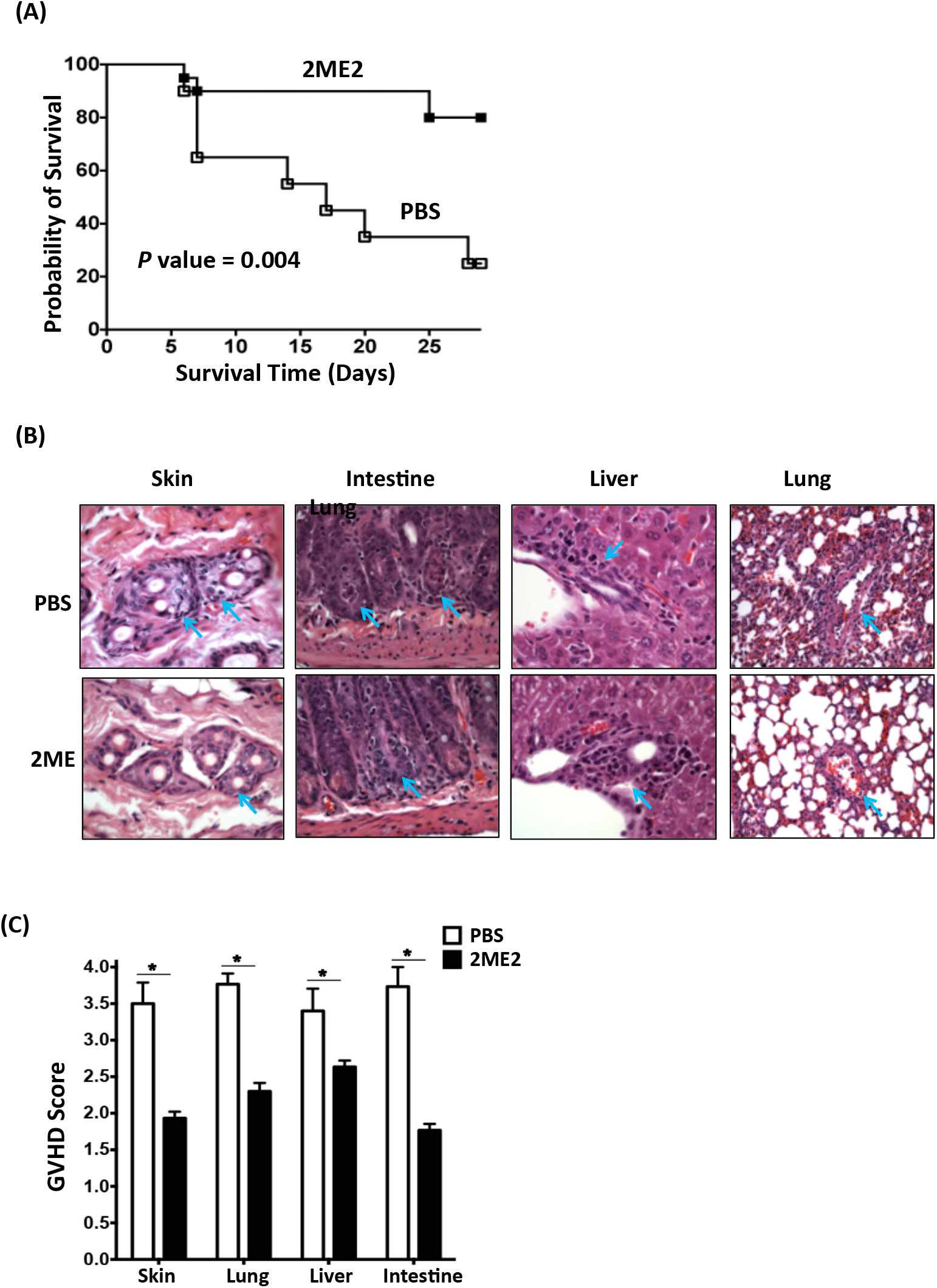
2ME2 treatment reduces mortality/morbidity associated with acute GVHD in mice. Lethally irradiated Balb/C mice received BM cells and splenocytes from C57BL/6 mice. Recipient mice were treated with 2ME2 (25 mg/kg) and PBS daily starting 24 hrs post-transplnt. (A) Survival was monitored and results were from 3 independent experiments (5 mice per group). Distributions of time to death were estimated using the Kaplan-Meier method, and groups were compared using the log-rank test. (B) Mice were sacrificed on day 7 post-transplant (5 mice per group). Tissues were placed in 10% formalin, embedded in paraffin, sectioned, and stained with hematoxylin and eosin, and scored for GVHD histopathology. The top and lower panels are representative sections from the skin, small intestine, liver and lung of mice treated with PBS control and 2ME2 respectively. Arrows indicate the histopathological changes in the skin, intestine, liver and lung. (C) The average score of skin, small intestine, liver and lung of each group. Results are shown as mean ± S.D. (t test; n=4 in each group). Asterisk represents data with *p* value less than 0.05 in t test.

### 2ME2 inhibits donor T cell proliferation in peripheral lymph node and Peyer’s patches

Donor T cells activation in the secondary lymphoid organs is a critical step in the development of acute GVHD. As shown in Figure 2A, we analyzed donor-derived T cells in 2ME2-treated mice 7 days post-transplant, and found a significantly lower number of donor-derived T cells (CD3^+^/H-2D^b+^) in the lymph node (LN) and Peyer’s patches (PP) of 2ME2-treated mice compared to that of control recipients (Figure 2A). However, the total number of donor-derived T cells in the spleen did not decrease. Further analysis demonstrated that 2ME2 treatment reduced both donor-derived CD4^+^ and CD8^+^ subsets in LN and PP. 2ME2 is known to bind to tubulin at the colchicine-binding site and disrupt tubulin, which resulting in the arrest of G2/M and apoptosis in dividing cells while sparing quiescent cells^11-12^. Using MLR, we investigate whether the reduction of donor-derived T cells observed in the secondary lymphoid organs is due to the inhibition of naïve T cells proliferation and cell-cycle arrest in the presence of 2ME2. As shown in Figure 2B, both CD4^+^ and CD8^+^ T cell proliferation was greatly reduced in the presence of 2ME2 at the 1-4 μM concentration range. At the concentration of 4 μM, the frequency of dividing cells was significantly lower in the 2ME2-treated samples compared to none-treatment control (2.8% vs. 30.4% in CD4^+^ cells; 0.15% vs. 9.8% in CD8^+^ cells). We further investigated whether 2ME2 plays a role in cell-cycle progression and mitosis in naïve T cell proliferation. As shown in Figure 2C, cells in the G1 and G2/M phases were greatly reduced and it was dose-dependent. Collectively, 2ME2 treatment at low dose (25 mk/kg) could affect donor T cell proliferation in the peripheral lymph node and Peyer’s patches in vivo, and thus contribute to the GVHD protection.

**Figure 2.**
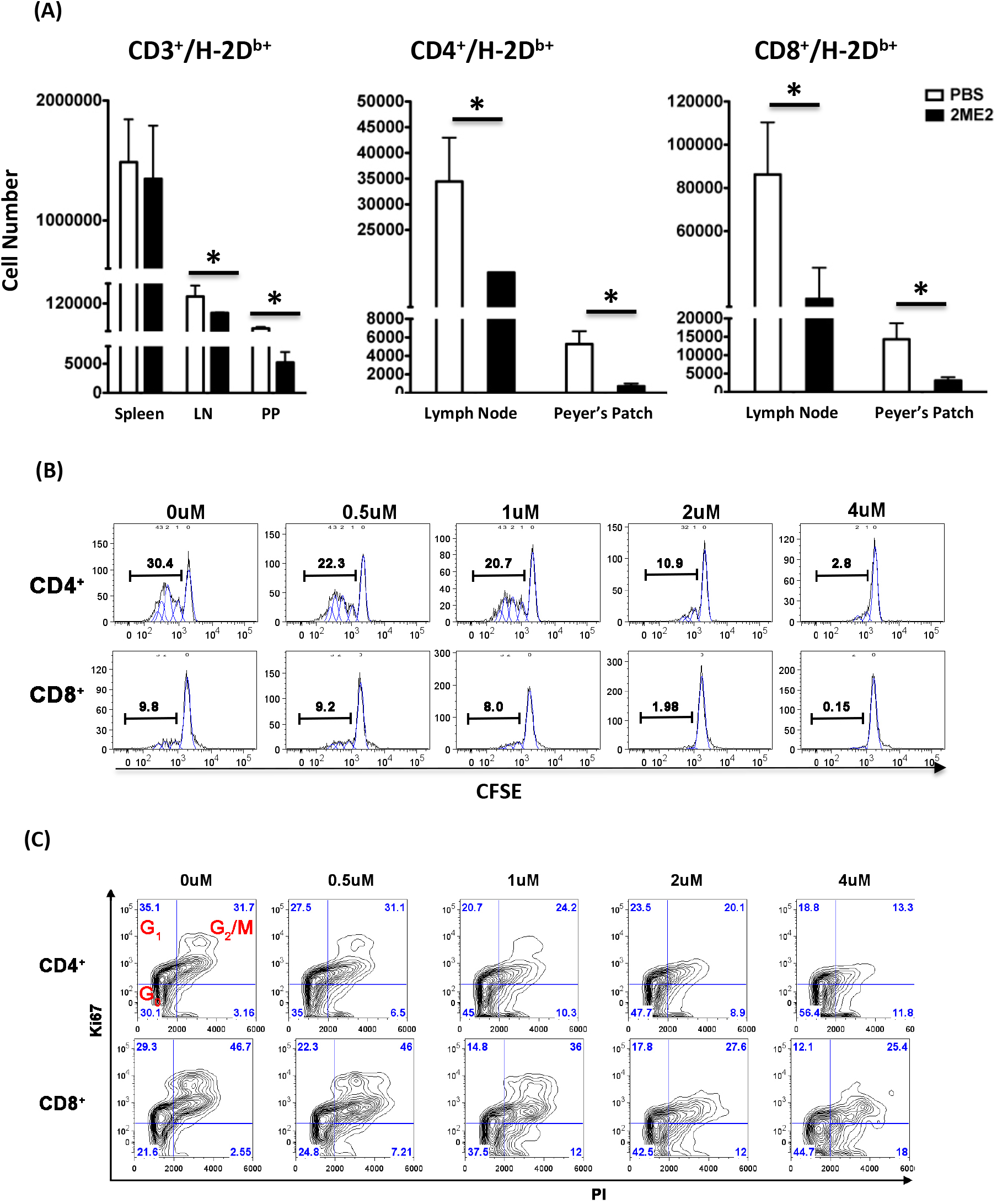
2ME2 inhibits the proliferation of mouse T cells. (A) Donor-derived T cells in the secondary lymphoid organs. Cells were harvested from mice on day 7 post-transplant. The number of donor-derived T cells (H-2D^b+^), CD4^+^H-2D^b+^ and CD8^+^H-2D^b+^ subsets were determined by FACS. Results are shown as bar graphs representing means ± S.D. (* is for *P* < 0.05). LN: lymph node; PP: Peyer’s patches. (B, C) Mouse T cell proliferation (B) and cell cycle progression (C) in MLR. Responder cells form C57BL/6 mice were plated at 1×10^6^ cells/ml in a volume of 200 µl/well and cocultured at a ratio of 2:1 with 3400 cGy irradiated stimulator cells from the Balb/C mice, in the presence of various concentration of 2ME2 as indicated. Proliferation was assessed on day 5 by CFSE dye dilution after sequential gating on lymphocytes (by scatter) and CD4^+^ or CD8^+^ T cells, the representative data of 3 independent samples were presented (B). For cell cycle analyses, cells were fixed and permeabilized, then stained with CD4 or CD8, Ki-67 and PI. Cells in the G0, G1 and G2/M phases of the cell cycle were determined by double staining for expression of Ki-67 and DNA content (PI) as indicated (C). Results shown are representative data from three independent experiments.

### Inhibition of inflammatory cytokine production upon 2ME2 treatment

Recently, it has been reported that 2ME2 modulates T cell cytokine production in EAE mouse model^21^. We further investigated the functional consequences of 2ME2 treatment on inflammatory cytokine production in vitro and in GVHD mice. In the presence of 2ME2 at various concentrations (0.5-2 uM), the frequency of both IFNγ^+^ and IL-17^+^ CD4^+^ cells was reduced upon allo-stimulation (Figure 3A). It was reported that the ability of 2ME2 to inhibit IL-17 production in Th17-polarized human T cells in vitro, and we found 2ME2 treatments can also reduce IL-17-producing CD4^+^ T cells in mouse MLR. Furthermore, the percentage of IFNγ^+^ subset of donor-derived CD4^+^ T cells was reduced in the spleen (26.5% vs 31.2%) of 2ME2-treated mice compared to the control recipients (Figure 3B). The same trend was observed for IL-2^+^ subset of donor-derived CD4^+^ cells (26.5% vs 31.2%). However, the percentage of donor-derived IL-17^+^CD4^+^ cells in the spleen of 2ME2-treatment mice did not decrease in comparison of control recipients. In addition, the reduction of IFN-γ and IL-2 was also observed in the serum collected 7 days post-transplant in these mice, whereas the amount of IL-17 remained the same in the serum of both 2ME2-treated and control mice (Figure 3C). The results demonstrated that 2ME2 treatment inhibitsTh1 not Th17 polarization, and reduces IL-2 production in GVHD mice.

**Figure 3.**
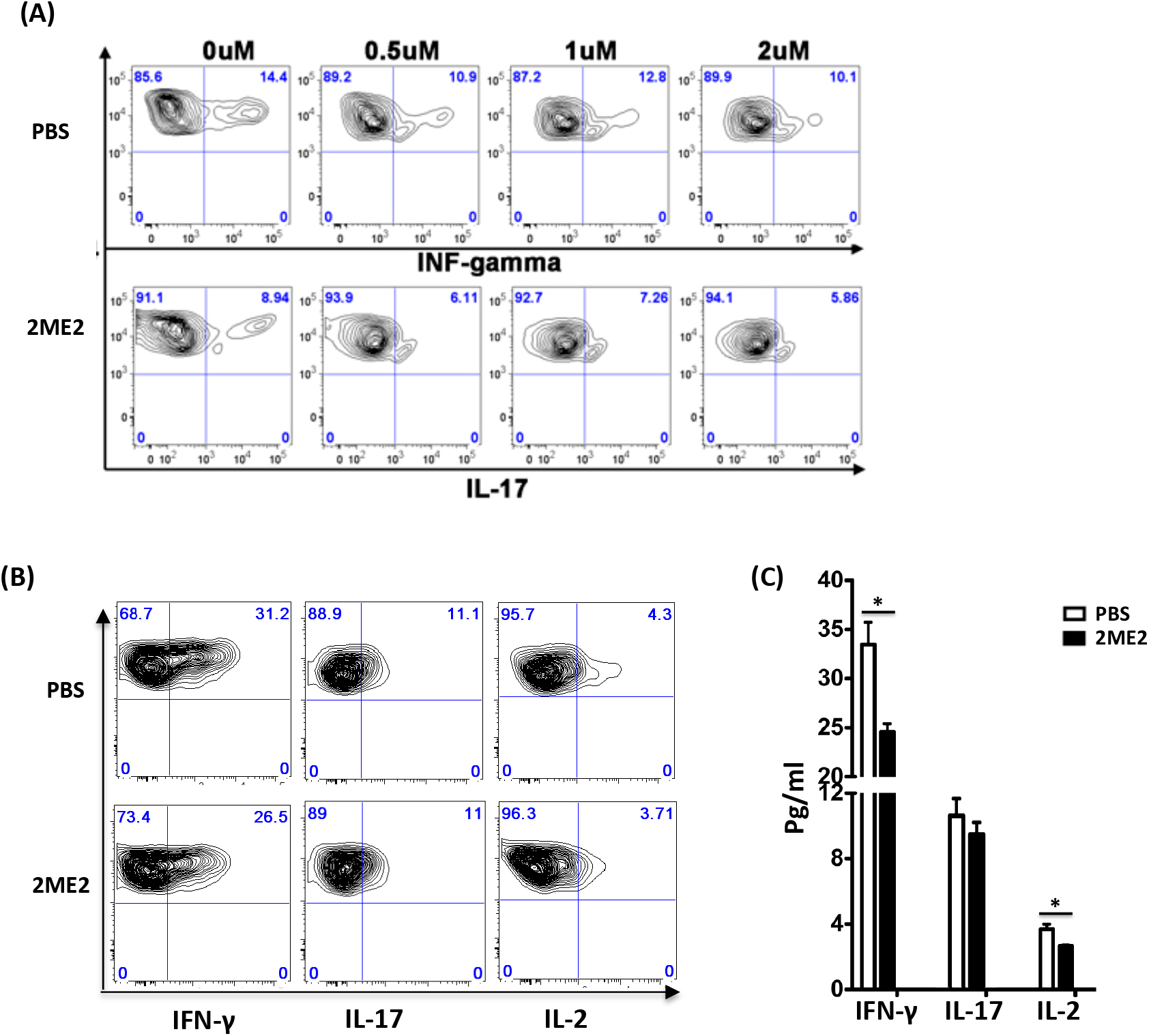
The differentiation and polarization of mouse CD4^+^ T cells in the presence of 2ME2. Activated mouse T cells were collected from in vitro MLR as described (A), or from spleen 7 days post-transplant (B and C). Cells were stimulated with PMA and ionomycin in the presence of Golgi-stop for 5 hrs, and stained with antibodies. The percentage of IL-17^+^, IFNγ^+^ and IL-2^+^ cells from donor-derived CD4^+^ cells (CD4^+^H-2D^b+^) was determined by FACS from four independent experiments. The data were the representative contour-plots of intracellular cytokine staining gated on CD4^+^H-2D^b+^ cells in vitro (A) and in vivo (B). Results are shown as bar graphs representing means ± S.D. (* is for P < 0.05) (C).

### 2ME2 treatment has no effect on the differentiation and cytolytic activity of CD8^+^ effector T cells

We further examined donor CD8^+^ T cell differentiation in the spleen of 2ME2-treated mice. Mouse CD8^+^ T cells were classified into three distinct subsets based on CD44 and CD62L expression: a naïve population, which is CD44^-^CD62L^+^; a central memory population, which is CD44^+^CD62L^+^; an effector population, which is CD44^+^CD62L^-28-29^. As shown in Figure 4A, the activated CD8^+^ T cell compartment from the PBS control spleen consisted of 5.33% naïve (N), 6.67% central memory (CM) and 86.5% effector (E) donor-derived CD8^+^ T cells on 7 day post-transplant. 2ME2 treatment did not alter the activation and differentiation of donor-derived CD8^+^ T cells in the spleen, and 94.2% of that differentiated into effector subset (CD44^+^CD62L^-^) in the spleen, which was comparable to the control mice. Furthermore, the expression level of CD107a on the donor-derived CD8^+^ T cells isolated from spleen of 2ME2-treated mice was similar as that of PBS control mice, indicating the cytoxic function was intact (Figure 4B). To determine whether 2ME2 treatment has any effect on CTL responses, we directly assessed the cytolytic activity of T cells in vitro (Figure 4C). CD8^+^ T cells were purified from C57/B6 mice and activated with irradiated APCs from Balb/C mice in vitro. For target cells, we used mouse leukemia cell line, P815 (H-2^d^, mastocytoma derived from a DBA/2 mouse). We found that effector T cells can specifically lysis target cells in the presence of 2ME2 at various concentrations (1-8 uM), and the cytolytic activity was similar as the control. These results indicate that 2ME2 treatment does not impair the cytotoxic activity of CD8^+^ effectot T cells.

**Figure 4.**
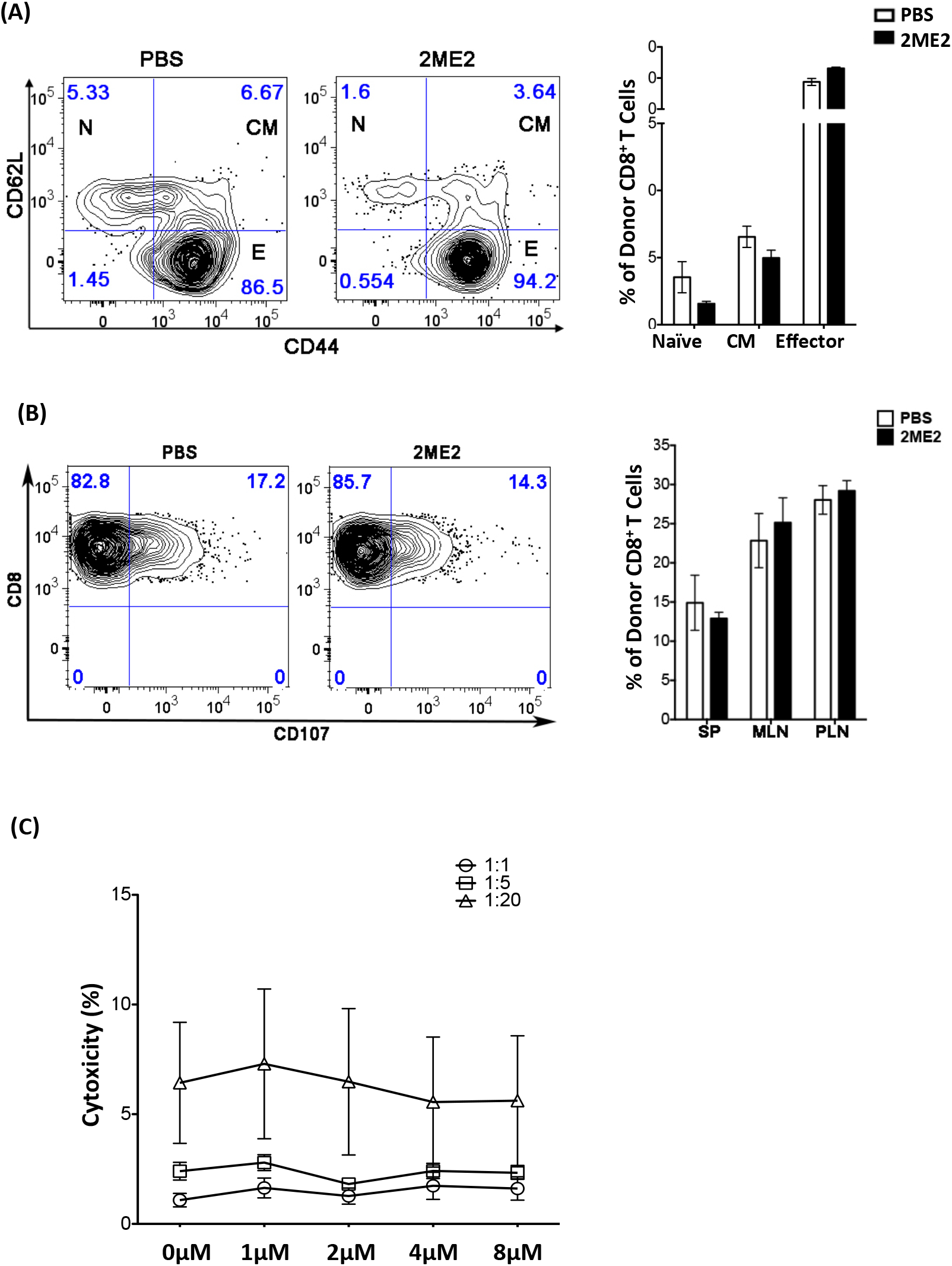
2ME2 has no effect on CD8^+^ T cell differentiation and cytoxicity: (A, B) Cells from spleen were collected 7 days post-transplant. A panel of surface antibodies were used to determine naïve, central memory and effector subset. The data were the representative contour-plots of CD44 and CD62L staining gated on CD8^+^H-2D^b+^ cells (A), or CD8 and CD107a staining gated on H-2D^b+^ cells (B). Results are shown as bar graphs representing means ± S.D. (* is for P < 0.05). (C) Cytotoxic activity of CD8^+^ effector T cells. The P815 (H-2^d^) target cells were labeled with DDAO-SE and incubated with in vitro–primed CD8^+^ effector cells from C57BL/6 mice at the E:T ratios indicated in the presence of 2ME2. Cells were cocultured for 4 hours before specific lysis was determined by staining with a phycoerythrin-conjugated anticleaved caspase-3 rabbit mAb (BD Biosciences). The stained cells were analyzed using a BD FACScanto II flow cytometer. Data shown here are representative of three independent experiments.

### 2ME2 treatment can preserve GVL activity post-BMT

To investigate whether GVL activity was maintained in the presence of 2ME2 treatment, we used P815 leukemia cell line, which migrates to the liver and bone marrow with infiltration in secondary lymphoid organs^27^. The P815 cells were transduced with a lentivirus vector coding for firefly luciferase (FLuc) for bioluminescence imaging. As shown in Figure 5A, the serial BLI images showed leukemic infiltration on day 7 after transplantation in most animals demonstrating similar leukemic engraftment (dorsal view only). However the rapid tumor growth was seen in mice received P815 and bone marrow cells alone (P815+BM) with PBS or 2ME2 treatment, as illustrated by the photon flux for mice on day 10, 14 and 19 post-transplant (Figure 5B).

**Figure 5.**
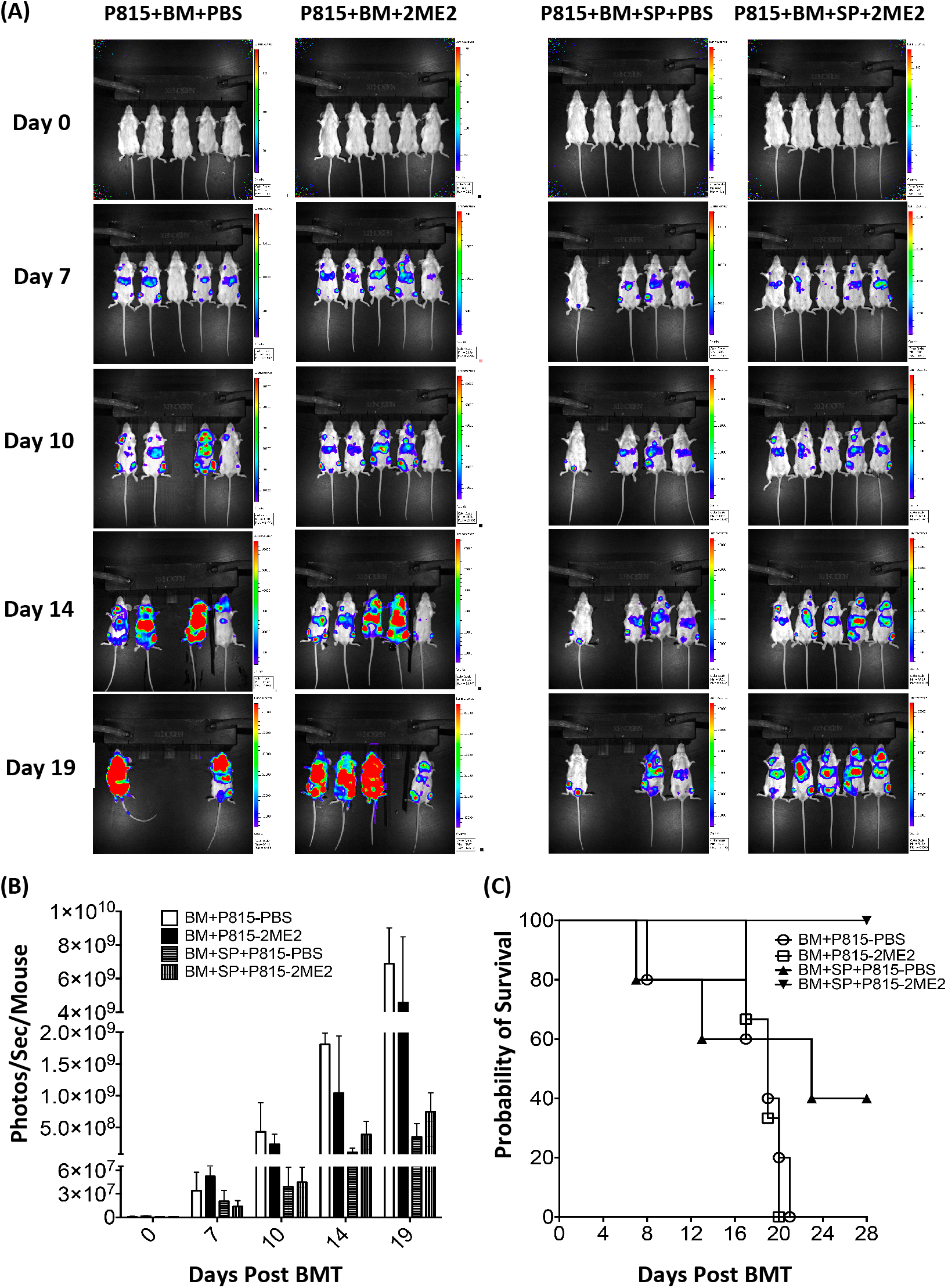
GVL function of is preserved in the presence of 2ME2 treatment. Recipient Balb/c mice received P815-Fluc on the same day of transplant as described. (A) Representative in vivo BLI of P815 leukemia cell-bearing mice with and without 2ME2 treatment in the presence or absence of donor T cells. Distribution of P815-Fluc in all surviving mice from a single experiment is displayed on days 0, 7, 10, 14 and 19 post-transplant. The experiment was performed twice with 5 mice per group and showed similar results. (B) Quantification of dorsal view BLI data. Photons emitted from the P815-Fluc cells in vivo are shown over time for the indicated groups. (C) Survival of Balb/c recipients post-transplant. Results were from two independent experiments (5 mice per group). BM: bone marrow cells; SP: splenocytes.

Images and photon flux showed P815 leukemic engraftment with significantly reduced signal and leukemia infiltration in mice received P815, bone marrow and splenocytes (P815+BM+SP) with or without 2ME2 treatment (Figure 5A and 5B). The overall survival of P815-bearing mice was prolonged in the mice received P815+BM+SP, in comparison to mice received P815+BM alone (Figure 5C). The data demonstrated 2ME2 treatment appears to preserve the GVL activity against P815 in mouse model.

## DISCUSSION

GVHD is the result of alloimmune responses elicited by donor T cell activation, proliferation and production of inflammatory cytokines^3, 6-10^. The same alloimmune responses are strongly associated with the beneficial GVL effect, and the molecular and cellular basis are not well understood^1^. It has been reported that exogenous 2ME2 prevents cell cycle progression and induces apoptosis in rapidly dividing cells^11-12^. Recently, 2ME2 has been shown to have anti-inflammatory activities in autoimmune diseases by modulating T cell polarization and cytokine production^18-22^. Our data demonstrate that 2ME2 treatment can reduce GVHD mortality and morbidity by inhibiting donor T cell proliferation, reduce the production of IFN-γ and IL-2 in donor-derived CD4^+^ T cells. Moreover, the cytolytic function of CTL is not impaired in vitro and the GVL activity remains largely intact in vivo in the presence of 2ME2.

Current data indicates that GVHD is a systemic immune reaction with a Th1 CD4 response, CD8 cytotoxic T lymphocyte (CTL) generation and an inflammatory cytokine storm^3^. Recent data investigated extensively CD4^+^ T cell subsets as essential regulators of adaptive immune responses and GVHD^8^. Various approaches to modify T cell mediated alloreactivity, and immune modulator agents are being explored for novel GVHD therapeutics^9-10^. Our data demonstrated that 2ME2 treatment at 25 mg/kg can protect mice from GVHD related mortality (Figure 1A). 2ME2 is known to induce cell cycle arrest and apoptosis in rapidly dividing cancer cells in patients at a dose proximately equivalent to 200mg/kg in mice^29^. We found there was a significant reduction of donor-derived CD4^+^ and CD8^+^ T cell in the peripheral lymph node and Peyer’s patches, where they were activated by the antigen-presenting cells (APCs) in GVHD initiation phase (Figure 2A). The reduction of donor-derived T cells observed in these secondary lymphoid organs is probably due to the antimitogenic properties of 2ME2. Specifically, it has been demonstrated that a significant decrease of naïve T cells proliferation (Figure 2B) and cell-cycle arrest (Figure 2C) upon 2ME2 treatment in MLR. Moreover, 2ME2 treatment does not impact engraftment and hematopoiesis, with granulocytes and monocytes found to be all donor-derived 3 weeks post-transplant (data not shown).

Recent published studies also found Th1/17 polarization and regulatory T cell (Treg) regulate GVHD in mice and human^8^. Spleen is the major site of donor T cell activation in GVHD, and the total number of donor-derived T cells was not decreased with low dose 2ME2 treatment (Figure 2A).

Although donor T cell proliferation is largely intact in the spleen, 2ME2 treatment can inhibit Th1 polarization and reduce IL-2 production in CD4^+^ T cells in GVHD mice (Figure 3B). The anti-inflammatory activity of 2ME2 in autoimmune diseases has been attributed to its ability to inhibit Th17 polarization^21-22^. Indeed, we found that 2ME2 treatment reduces both Th1 (IFNγ-producing) and Th17 (IL-17-producing) differentiation in MLR (Figure 3A). However, the percentage of donor-derived IL17^+^CD4^+^ cells in spleen and IL-17 level in the serum of 2ME2-treatment mice are similar to that of control recipients. We conclude that the reduced GVHD mortality and morbidity in low dose 2ME2-treated mice is associated with the decrease of donor Th1 not Th17 polarization.

The same T cell-mediated alloimmune responses in GVHD are strongly associated with the beneficial graft-versus leukemia (GVL) effect, and the molecular and cellular basis are not well understood^1^. Previous reports demonstrate that the proliferation and differentiation of CD4^+^ T cells are responsible for GVHD while CD8^+^ effector T cells contribute to GVL ^1, 3, 7-10^. We found that low dose 2ME2 treatment has no effect on the proliferation and differentiation of CD8^+^ effector T cells in the spleen of GVHD mice (Figure 4A), and does not impair the cytotoxic activity of CD8^+^ effector T cells in vitro (Figure 4C). Furthermore, 2ME2 treatment can minimize GVHD without compromising the GVL effect in vivo (Figure 5). It is possible that the treatment with 2ME2 inhibits leukemia growth independent of T-cell–mediated GVL effect. However, we observed that 2ME2 treatment at current dose and schedule did not affect P815 growth in the recipients with BM alone (Figure 5A). Taken together, it is clear that 2ME2 treatment impairs donor T-cell responses, which contributes to GVHD while sparing GVL effects mediated by cytotoxic T cells.

Our studies have established the disease-modifying activity of 2ME2 in mouse GVHD model, and thus raise the possibility for using this drug as a potential new therapeutic intervention. Although novel approaches are being constantly evaluated to minimizing GVHD without compromising the GVT effect, it is important to dissect the role of 2ME2 treatment in regulating T cell activation, demonstrate GVHD and GVL effect in mouse models. The study will not only significantly advance our knowledge of GVHD, but also provide a rationale and pre-clinical data for using 2ME2 as a novel therapy in patients.

## Acknowledgements

The work is supported in part by NIH/NIAID grant 1R21AI101932 (Q.M.), NIH/NCI grant 1P01CA148600 (Q.M. and J.J.M.). D.L. and R.P. performed research. P.Z. provided reagents. D.L., R.P., R.B. and Q.M. analyzed data. A.M.A., R.E.C., J.J.M. and T.W.M. participated in study design and discussions. Q.M. designed the research, analyzed data and wrote the paper. We have no conflicts of interest to disclose.

## REFERENCES

1. Appelbaum FR. 2001. Haematopoietic cell transplantation as immunotherapy. Nature. 411:385–9.

2. Sale, GE. The Pathology of Organ Transplantation. London, Stoneham: Butterworth Publishers, Inc. 1990. p.229–260.

3. Ferrara JLM, Antin JH. The Pathophysiology of graft-versus-host disease. In: Thomas ED, Blume KG, Forman SJ, eds. Hematopoietic Cell Transplantation. Blackwell Scientific Publications, Boston. 1994. p.305–15.

4. Alousi AM, Bolaños-Meade J, Lee SJ. 2013. Graft-versus-host disease: state of the science. Biol Blood Marrow Transplant. 1 Suppl:S102–8.

5. Levine JE, Logan B, Wu J, Alousi AM, Ho V, Bolaños-Meade J, Weisdorf D, Blood and Marrow Transplant Clinical Trials Network. 2010. Graft-versus-host disease treatment: predictors of survival. Biol Blood Marrow Transplant. 16(12):1693–9.

6. Fowler DH, and Gress RE. 1997. Graft-versus-host disease as a Th1-type process: regulation by donor cells of Th2 cytokine phenotype. In: Ferrara JLM, Deeg HJ, Burakoff SJ, eds. Graft-vs.-Host Disease. New York: Marcel Dekker Inc. p.479–500.

7. Shlomchik WD. 2007. Graft-versus-host disease. Nat Rev Immunol. 7:340–352.

8. Coghill JM, Sarantopoulos S, Moran TP, Murphy WJ, Blazar BR, Serody JS. 2011. Effector CD4+ T cells, the cytokines they generate, and GVHD: something old and something new. Blood. 117:3268–76.

9. Li HW, Sykes M. 2012. Emerging concepts in haematopoietic cell transplantation. Nat Rev Immunol. 12(6):403–16.

10. Blazar BR, Murphy WJ, Abedi M. Advances in graft-versus-host disease biology and therapy. 2012. Nat Rev Immunol. 12(6):443–58.

11. LaVallee TM, Zhan XH, Herbstritt CJ, Kough EC, Green SJ, Pribluda VS. 2002. 2-Methoxyestradiol inhibits proliferation and induces apoptosis independently of estrogen receptors alpha and beta. Cancer Res. 62(13):3691–7.

12. Pribluda VS, Gubish ER Jr, Lavallee TM, Treston A, Swartz GM, Green SJ. 2000. 2-Methoxyestradiol: an endogenous antiangiogenic and antiproliferative drug candidate. Cancer Metastasis Rev. 19(1-2):173–9.

13. Sutherland TE, Anderson RL, Hughes RA, Altmann E, Schuliga M, Ziogas J, Stewart AG. 2007. 2-Methoxyestradiol--a unique blend of activities generating a new class of anti-tumour/anti-inflammatory agents. Drug Discov Today. 12(13-14):577–84.

14. Berg D, Sonsalla R, Kuss E. 1983. Concentrations of 2-methoxyoestrogens in human serum measured by a heterologous immunoassay with an 125I-labelled ligand. Acta Endocrinol (Copenh). 103(2):282–8.

15. Huh JI, Qiu TH, Chandramouli GV, Charles R, Wiench M, Hager GL, Catena R, Calvo A, LaVallee TM, Desprez PY, Green JE. 2007. 2-methoxyestradiol induces mammary gland differentiation through amphiregulin-epithelial growth factor receptor-mediated signaling: molecular distinctions from the mammary gland of pregnant mice. Endocrinology. 148(3):1266–77.

16. Ostensen M, Villiger PM. 2007. The remission of rheumatoid arthritis during pregnancy. Semin Immunopathol. 29(2):185–91.

17. Offner H, Polanczyk M. 2006. A potential role for estrogen in experimental autoimmune encephalomyelitis and multiple sclerosis. Ann N Y Acad Sci. 1089:343–72.

18. Josefsson E, Tarkowski A. 1997. Suppression of type II collagen-induced arthritis by the endogenous estrogen metabolite 2-methoxyestradiol. Arthritis Rheum. 40(1):154–63

19. Plum SM, Park EJ, Strawn SJ, Moore EG, Sidor CF, Fogler WE. 2009. Disease modifying and antiangiogenic activity of 2-methoxyestradiol in a murine model of rheumatoid arthritis. BMC Musculoskelet Disord. 1;10:46.

20. Issekutz AC, Sapru K. 2008. Modulation of adjuvant arthritis in the rat by 2-methoxyestradiol: an effect independent of an anti-angiogenic action. Int Immunopharmacol. 8(5):708–16.

21. Duncan GS, Brenner D, Tusche MW, Brüstle A, Knobbe CB, Elia AJ, Mock T, Bray MR, Krammer PH, Mak TW. 2012. 2-Methoxyestradiol inhibits experimental autoimmune encephalomyelitis through suppression of immune cell activation. Proc Natl Acad Sci U S A. 109(51):21034–9.

22. Zepp J, Wu L, Li X. 2011. IL-17 receptor signaling and T helper 17-mediated autoimmune demyelinating disease. Trends Immunol. 32(5):232–9.

23. Wang Y, Li D, Jones D, Bassett R, Sale GE, Khalili J, Komanduri KV, Couriel DR, Champlin RE, Molldrem JJ, Ma Q. 2009. Blocking LFA-1 activation with lovastatin prevents graft-versus-host disease in mouse bone marrow transplantation. Biol Blood Marrow Transplant. 15:1513–22.

24. Li D, Molldrem JM, Ma Q. 2009. LFA-1 regulates CD8+ T cell activation via TCR-mediated and LFA-1-mediated Erk1/2 signal pathways. J Biol Chem. 284:21001-10.

25. Ma Q, Li D, Nurieva R, Patenia R, Bassett R, Cao W, Alekseev AM, He H, Molldrem JJ, Kroll MH, Champlin RE, Sale GE, Afshar-Kharghan V. 2012. Reduced GVHD in C3-deficient mice is associated with the decrease of donor Th1/Th17 differentiation. Biol Blood Marrow Transplant. 18:1174–81.

26. Li D, Li Y, Hernandez JA, Patenia R, Kim TK, Khalili J, Dougherty MC, Hanley PJ, Bollard CM, Komanduri KV, Hwu P, Champlin RE, Radvanyi LG, Molldrem JJ, Ma Q. 2010. Lovastatin prevents T cell proliferation without affecting the cytolytic function of EBV-, CMV- and MART1-specific CTLs. J Immunother. 33:975–82.

27. Hanash AM, Kappel LW, Yim NL, Nejat RA, Goldberg GL, Smith OM, Rao UK, Dykstra L, Na IK, Holland AM, Dudakov JA, Liu C, Murphy GF, Leonard WJ, Heller G, van den Brink MR. Abrogation of donor T-cell IL-21 signaling leads to tissue-specific modulation of immunity and separation of GVHD from GVL. Blood. 2011 Jul 14;118(2):446–55.

28. Dutt S, Baker J, Kohrt HE, Kambham N, Sanyal M, Negrin RS, Strober S. 2011. CD8+CD44(hi) but not CD4+CD44(hi) memory T cells mediate potent graft antilymphoma activity without GVHD. Blood. 117:3230–9.

29. Zheng H, Matte-Martone C, Jain D, McNiff J, Shlomchik WD. 2009. Central memory CD8+ T cells induce graft-versus-host disease and mediate graft-versus-leukemia. J Immunol. 182(10):5938–48.

30. Escuin D, Kline ER, Giannakakou P. 2005. Both microtubule-stabilizing and microtubule-destabilizing drugs inhibit hypoxia-inducible factor-1alpha accumulation and activity by disrupting microtubule function. Cancer Res. 65(19):9021–8.

